## Supplementary data for "Kin discrimination promotes horizontal gene transfer between unrelated strains in *Bacillus subtilis*"

**Supplementary Table 1 | Strains used in this study.**

| ***B. subtilis* strains** | **genotype** | **Source** |
| --- | --- | --- |
| PS-216 | Wild type | ^1^ |
| PS-196 | Wild type | ^1^ |
| PS-13 | Wild type | ^1^ |
| PS-218 | Wild type | ^1^ |
| PS-18 | Wild type | ^1^ |
| PS-68 | Wild type | ^1^ |
| 11A79 | *PcomGA-yfp (Cm)* | ^2^ |
| BD2121 | his leu met ΔcomK∷kan | ^3^ |
| BKK25750 | Δ*nucB::kan* trpC2 | ^4^ |
| BKE09190 | Δ*yhcR::erm* trpC2 | ^4^ |
| BKK24730 | Δ*comGA::kan* trpC2 (Kn) | ^4^ |
| BKE24730 | Δ*comGA::ery* trpC2 (Ery) | ^4^ |
| ZK4300 | Δ*epsA-O* (Tet) | ^5^ |
| NL362 | *sigW::erm* (Ery) | ^5^ |
| ZK4860 | *amyE*::P*sigW*-*yfp* (Sp) | ^5^ |
| BM1345 | PS-216 *amyE*::P43-*cfp* (Sp) | This work |
| BM1328 | PS-216 *sacA*::P43-*yfp* (Cm) | This work |
| BM1348 | PS-196 *amyE*::P43-*cfp* (Sp) | This work |
| BM1332 | PS-196 *sacA*::P43-*yfp* (Cm) | This work |
| BM1350 | PS-13 *amyE*::P43-*cfp* (Sp) | This work |
| BM1335 | PS-13 *sacA*::P43-*yfp* (Cm) | This work |
| BM1347 | PS-218 *amyE*::P43-*cfp* (Sp) | This work |
| BM1468 | PS-218 *sacA*::P43-*yfp* (Cm) | This work |
| BM1651 | PS-18 *amyE*::P43-*cfp* (Sp) | This work |
| BM1470 | PS-18 *sacA*::P43-*yfp* (Cm) | This work |
| BM1544 | PS-68 *amyE*::P43-*cfp* (Sp) | This work |
| BM1469 | PS-68 *sacA*::P43-*yfp* (Cm) | This work |
| BM1546 | PS-216 *amyE*::P43-*cfp* (Sp) P*comGA*-*yfp* (Cm) | This work |
| BM1655 | PS-216 *amyE*:: P43-*cfp* (Sp) Δ*comGA* (Kn) | This work |
| BM1583 | PS-216 *amyE*::P43-*cfp* (Sp) Δ*nucB* (Kn) Δ*yhcR* (Ery) | This work |
| BM1566 | PS- 196 *sacA*::P43-*yfp* (Cm) Δ*yhcR* (Ery) | This work |
| BM1556 | PS-216 *amyE*:: P43-*cfp* (Sp) Δ*comGA* (Ery) | This work |
| BM1577 | PS-216 *sacA*:: P43-*yfp* (Cm) Δ*comGA* (Ery) | This work |
| BM1578 | PS-196 *sacA*:: P43-*yfp* (Cm) Δ*comGA* (Ery) | This work |
| BM1657 | PS-196 *amyE*:: P43-*cfp* (Sp) Δ*comGA* (Ery) | This work |
| BM1698 | PS-218 *amyE*:: P43-*cfp* (Sp) Δ*comGA* (Kn) | This work |
| BM1703 | PS-218 *sacA*:: P43-*yfp* (Cm) Δ*comGA* (Ery) | This work |
| BM1699 | PS-13 *amyE*:: P43-*cfp* (Sp) Δ*comGA* (Kn) | This work |
| BM1636 | PS-13 *sacA*:: P43-*yfp* (Cm) Δ*comGA* (Ery) | This work |
| BM1070 | PS-216 Δ*epsA-O* (Tet) | This work |
| BM1666 | PS-196 Δ*epsA-O* (Tet) | This work |
| BM1418 | PS-216 Δ*QXP* (Kn) | This work |
| BM1646 | PS-216 *amyE*::P43-*cfp* (Sp) Δ*sigW* (Ery) | This work |
| BM1644 | PS-216 *sacA*::P43-*yfp* (Cm) Δ*sigW* (Ery) | This work |
| BM1647 | PS- 196 *amyE*::P43-*cfp* (Sp) Δ*sigW* (Ery) | This work |
| BM1645 | PS- 196 *sacA*::P43-*yfp* (Cm) Δ*sigW* (Ery) | This work |
| BM1869 | PS-218 *amyE*::P43-*cfp* (Sp) Δ*sigW* (Ery) | This work |
| BM1868 | PS-218 *sacA*::P43-*yfp* (Cm) Δ*sigW* (Ery) | This work |
| BM1847 | PS-13 *amyE*::P43-*cfp* (Sp) Δ*sigW* (Ery) | This work |
| BM1846 | PS-13 *sacA*::P43-*yfp* (Cm) Δ*sigW* (Ery) | This work |
| BM1642 | PS-216 *amyE*::P*sigW*-*yfp* (Sp) | This work |
| BM1643 | PS-196 *amyE*::P*sigW*-*yfp* (Sp) | This work |
| BM1871 | PS-13 *amyE*::P*sigW*-*yfp* (Sp) | This work |
| BM1872 | PS-18 *amyE*::P*sigW*-*yfp* (Sp) | This work |
| BM1873 | PS-68 *amyE*::P*sigW*-*yfp* (Sp) | This work |
| BM1874 | PS-218 *amyE*::P*sigW*-*yfp* (Sp) | This work |
| **Plasmids/*E.coli*** |  |  |
| pEM1069 | DH5α *amyE*::P43-*cfp* (Sp), Amp | This work |
| pEM1071 | DH5α *sacA*::P43-*yfp* (Cm), Amp | This work |
| Pkm8 | *amyE*:: *spoIIQ* -*cfp* (Sp), Amp | ^6^ |
| ECE174 | DH5a(p*Sac*-Cm) | ^7^ |
| pED302 | *comQXP*::Kn | ^8^ |

**Supplementary Table 2 |** **Strains combinations used in experiments.**

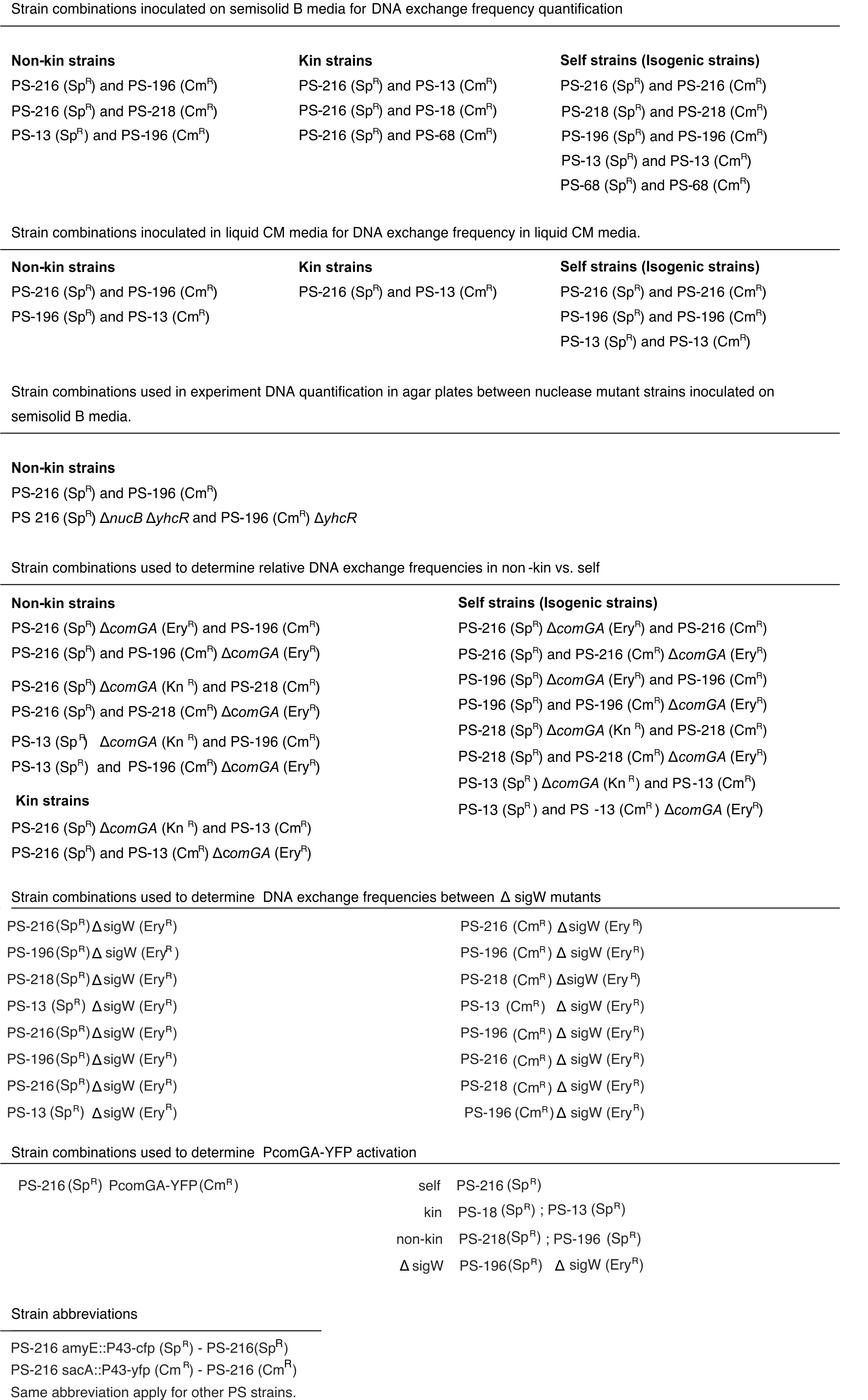

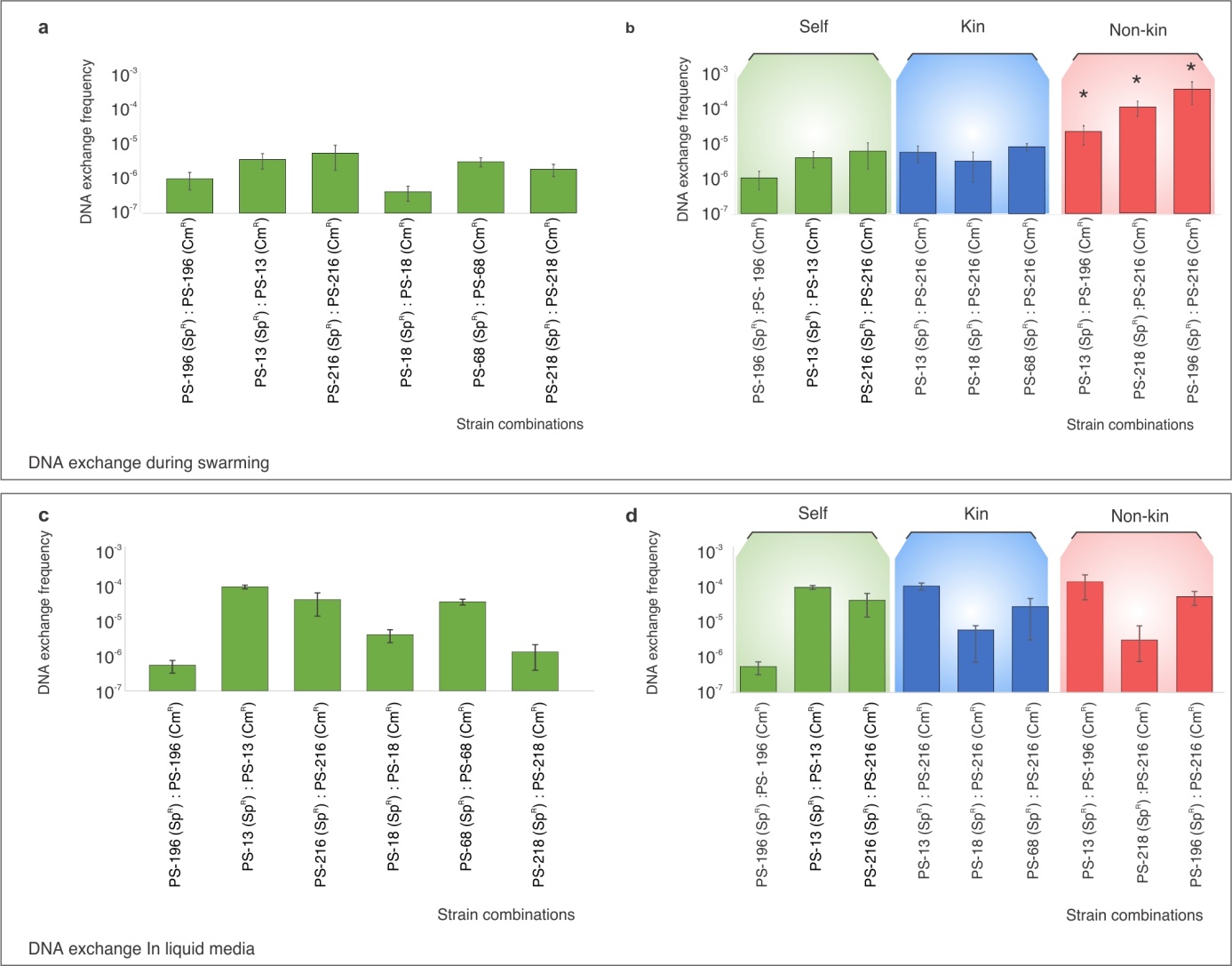

**Supplementary Fig. 1 | DNA exchange in liquid co-cultures and during swarming of self-controls and of strains with inverse antibiotic (Ab) markers. a,** DNA exchange between isogenic self-strains during swarming. Strain combinations tested were: PS-196Sp: PS-196Cm, PS-13Sp: PS-13Cm, PS-216Sp: PS-216Cm, PS-18Sp:PS-18Cm, PS-68Sp:PS-68Cm, PS-218Sp:PS-218Cm . **b**, DNA exchange of strains during swarming with inverse Ab and fluorescent markers. Strain combinations tested were: 196Sp:PS-196Cm, PS-13Sp:PS-13Cm, PS-216Sp:PS-216Cm, PS-13Sp:PS-216Cm, PS-18Sp:PS-216Cm, PS-68Sp:PS-216Cm, PS-13Cm:PS-196Sp, PS-218Sp:PS-216Cm, PS-196Sp:PS-216Cm. **c**, DNA exchange of self-self controls in liquid CM media. Strain combinations from left to right: PS-68Sp:PS-68Cm, PS-18Sp:PS-18Cm, PS-218Sp:PS-218Cm, PS-216Sp:PS-216Cm, PS-196 Sp:PS-196Cm, PS-13 Sp:PS-13Cm. **d**, DNA exchange in liquid co-cultures. Strains combinations from left to right: PS-196Sp:PS-196Cm, PS-13Sp:PS-13Cm, PS-216Sp:PS-216Cm, PS-13Sp:PS-216Cm, PS-18Sp:PS-216Cm, PS-68Sp:PS-216Cm, PS-68Sp:PS-216Cm, PS-13Sp:PS-196Cm, PS-218Sp:PS-216Cm, PS-196Sp:PS-216Cm. Strain abbreviations are as follows PS-216Sp (amyE::P43-cfp (SpR) and PS-216Cm (sacA::P43-yfp (CmR). Strains abbreviations are as follows: Sp - amyE::P43-cfp (SpR) and Cm - sacA::P43-yfp (CmR). All experiments were performed in three independent experiments using three replicates. Error bars represent SD of the average values. * represent statistically significant values (two tailed t-test, see Supplementary Information for details).

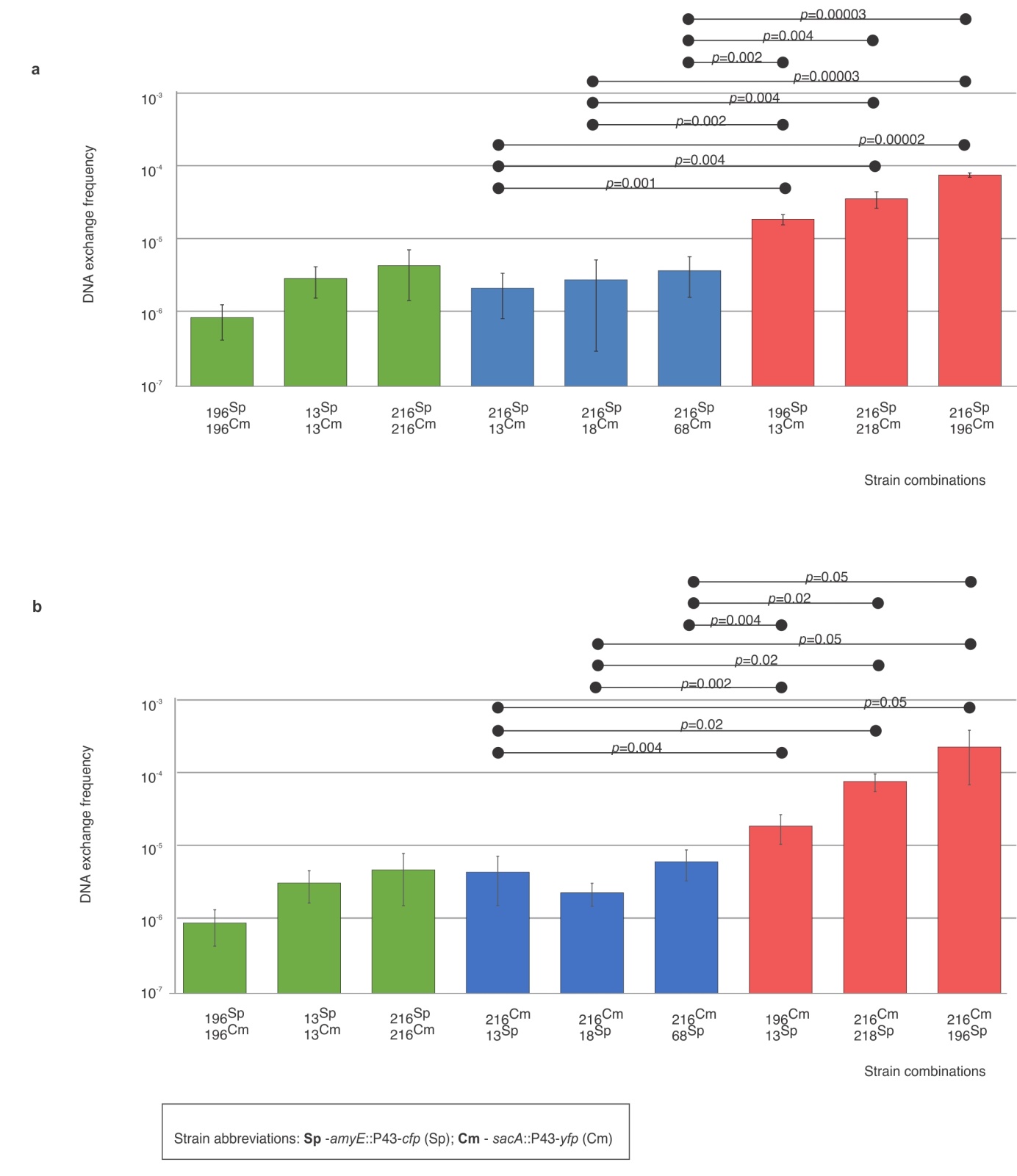

**Supplementary Fig. 2 | Non-kin DNA exchange differs from kin-DNA exchange.** **a**, DNA exchange frequency of kin and non-kin strains with the emphasis on *p* value (two tailed T-test) shown between experimentally obtained DNA exchange frequencies of kin sets compared to non-kin sets. **b**, DNA exchange frequency of kin and non-kin strains with inverse antibiotic markers with the emphasis on p value (two tailed t-test) between experimentally obtained figures of kin sets compared to non-kin sets. All experiments were performed in at least three replicates in three independent experiments. Error bars represent SD of the average values.

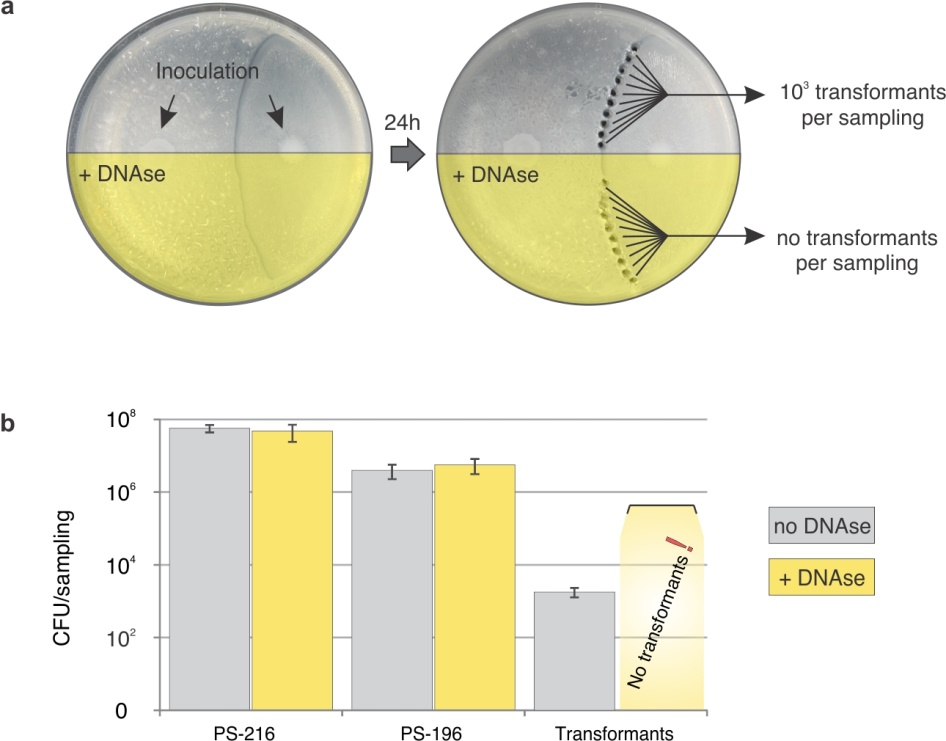

**Supplementary Fig. 3 | Extracellular DNA is required for DNA exchange. a**, Strain pairs were inoculated on agar plates of which half of each surface was covered with DNaseI (yellow area), and cells were harvested from the swarm boundary over both halves of the agar plate to quantify double recombinants (left) and total CFU counts. **b**, total CFU counts of each strain in the non-treated section (grey) and DNAseI treated section (yellow). Numerous transformants were obtained from the non-treated section of the agar plate, but no transformants could be recovered from the DNAse treated portion of the agar plate. All experiments were performed in three independent experiments using three replicates. Error bars represent SD of the average values.

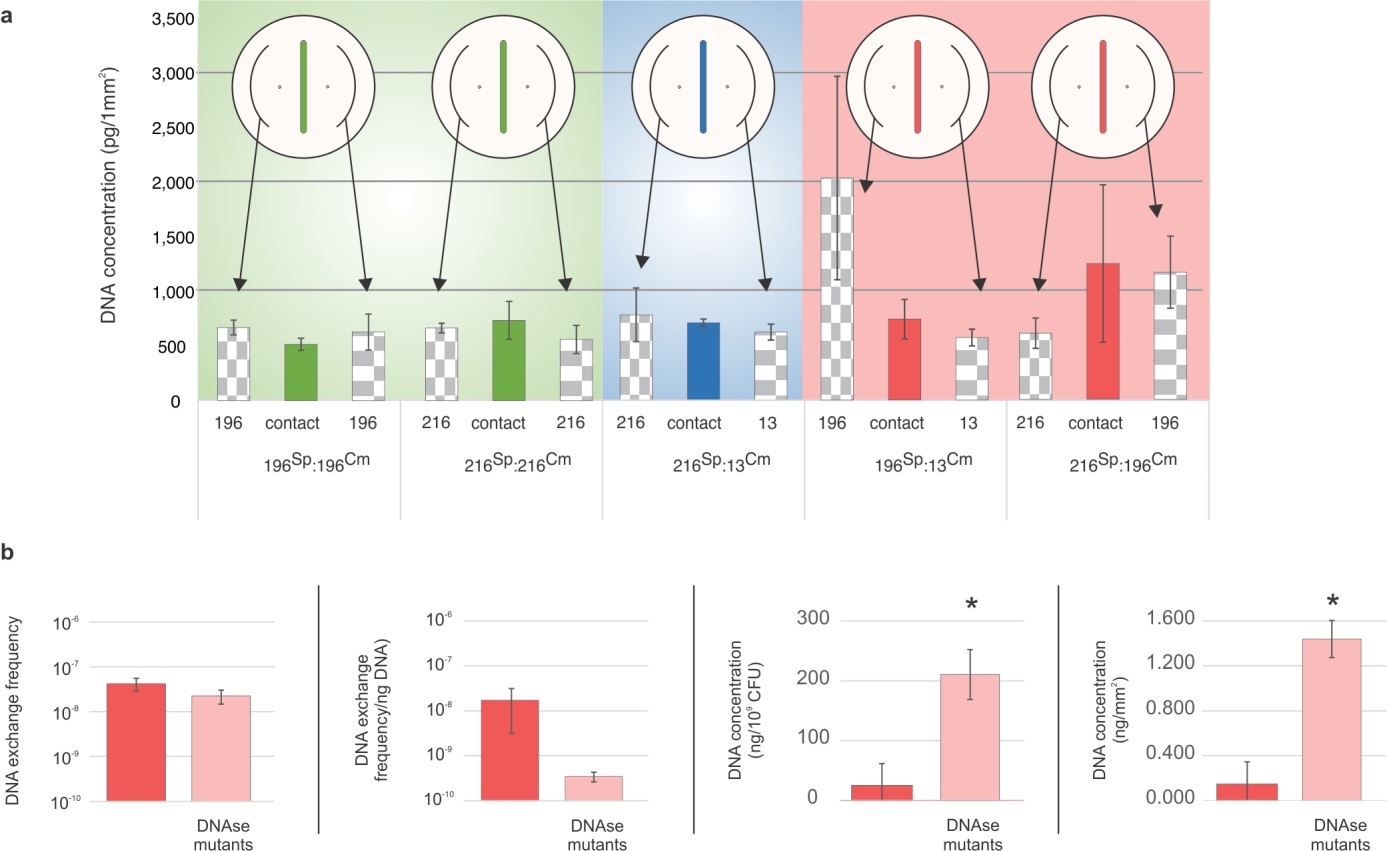

Supplementary Fig. 4. DNA concentration at the boundary. a, DNA concentration at the meeting points of self/kin (green), kin (blue) and non-kin strains (red). DNA concentration in the swarms is shown in grey columns. See table 2 for strain abbreviations. b, DNA exchange frequency and DNA concentration at the boundary between non-kin wt and between nuclease mutants (ΔyhcR and ΔnucB). Samples were taken from meeting points of two wild type strains (red) (PS-216 and PS-196) and two DNAse mutants (pink)(PS-216, ΔyhcR, ΔnucB and PS-196 ΔyhcR). From left to right: Transformation frequency between wt and between nuclease mutant strains, transformation frequency per ng DNA available at the meeting point, DNA concentration per cell (10^9^ CFU), DNA concentration per mm2 (ng). All experiments were performed in at least three replicates in three independent experiments. Error bars represent SD of the average values. * represent statistically significant values compared to corresponding wt pairings (two tailed t-test). For statistical parameters see Supplementary Results.

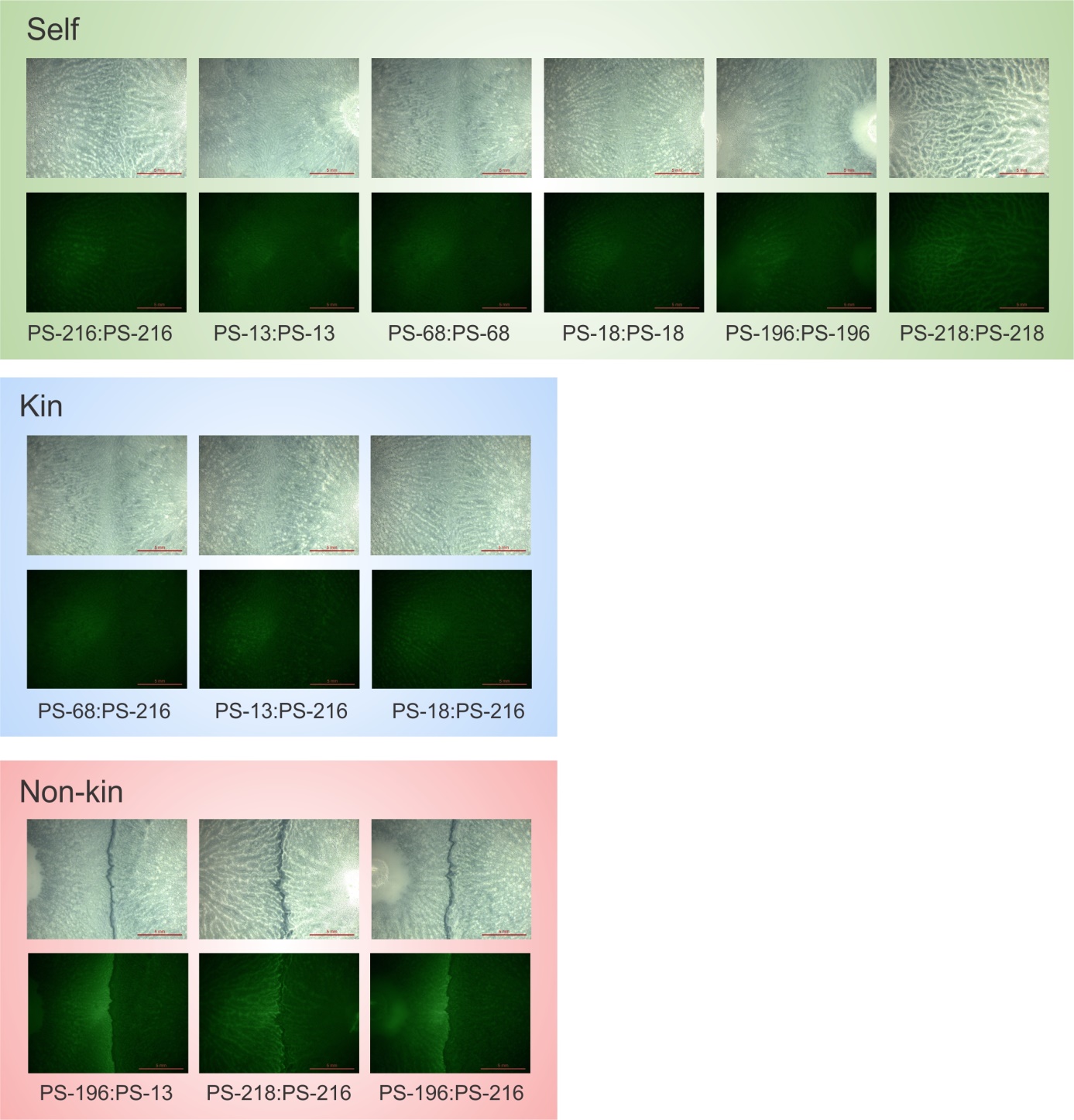

**Supplementary Fig. 5.** Stress and competence induction. Representative photos of meeting points of self (green, top) kin (blue, middle) and non-kin (red, bottom) *sigW-yfp* swarms. Both strains in the interaction were carrying the sigW-yfp.

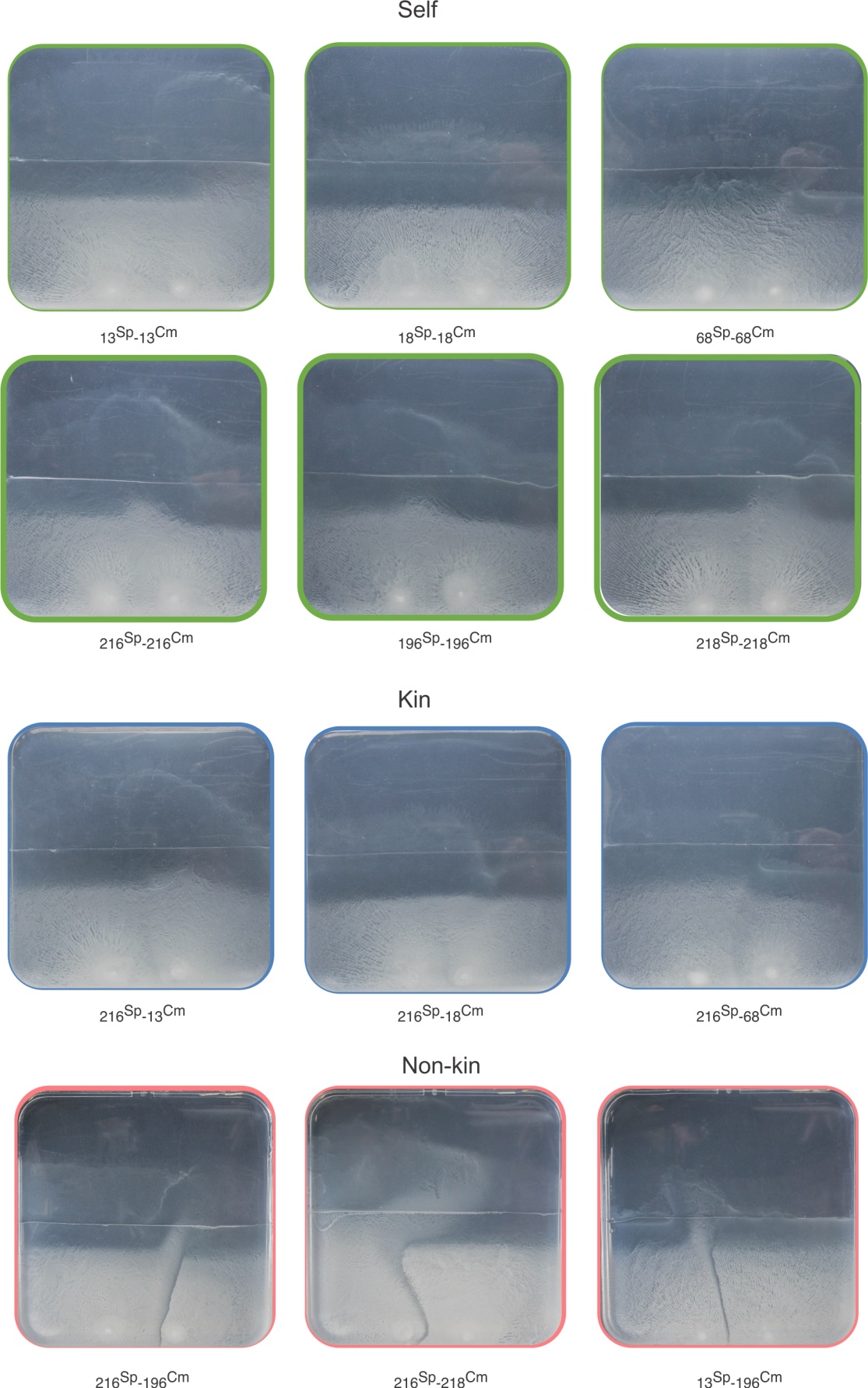

**Supplementary Fig. 6.** KD-mediated transformation can enable adaptation to novel selective pressures. Representative photos of meeting points of self (green, top) kin (blue, middle) and non-kin (red, bottom) strains.

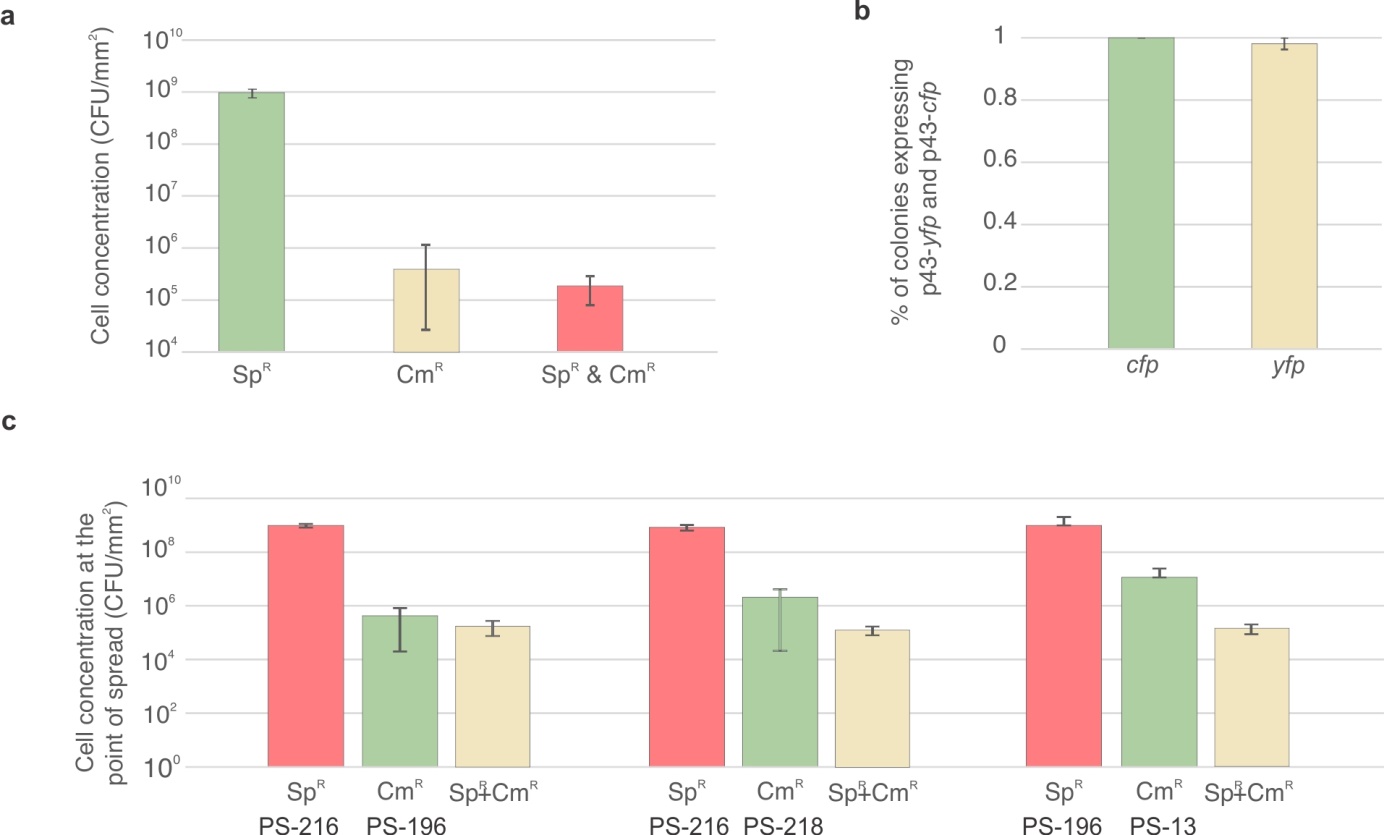

**Supplementary Fig. 7. DNA provides ecological advantage.** **a,** CFU of PS-216, PS-196 and transformants at the sampling points. **b,** The ratio of colonies expressing both *cfp* and *yfp* fluorescent proteins, isolated from the dual Ab sampling area. **c**) CFU of interacting non-kin strains carrying Cm, Sp or dual resistance genes at the sampling points. All experiments were performed in three independent experiments. Error bars represent SD of the average values.

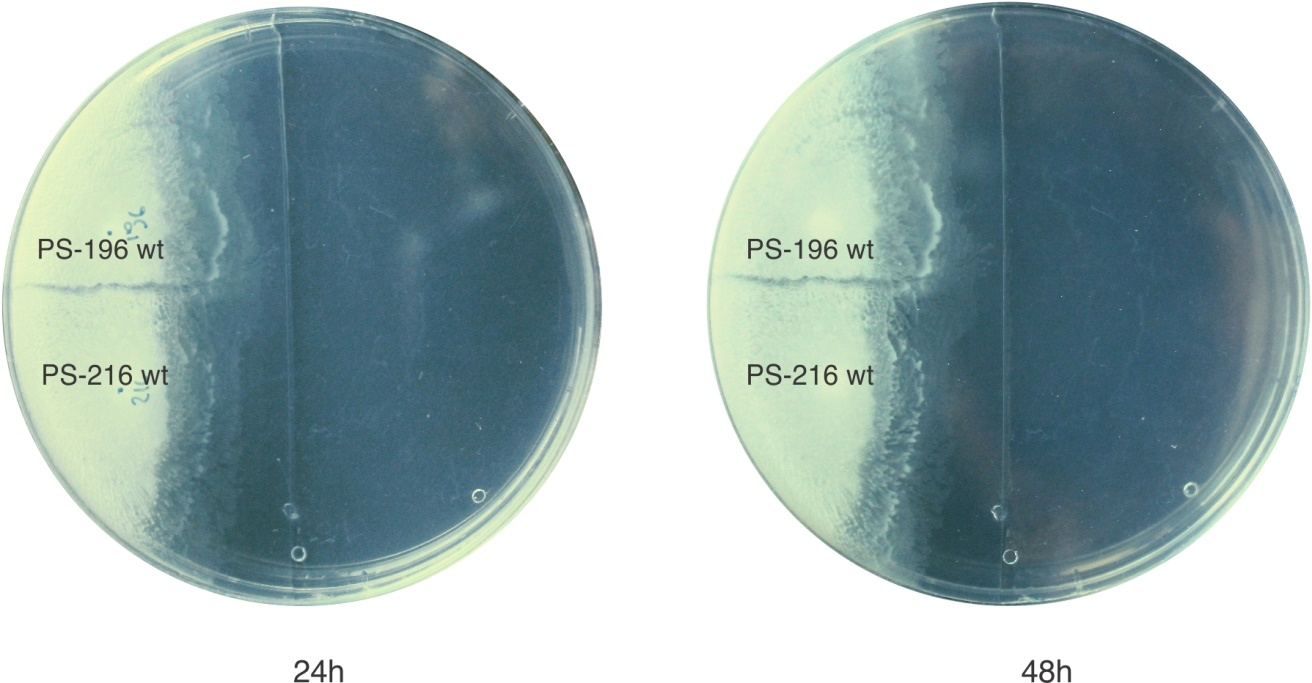

**Supplementary Fig. 8 | No spontaneous mutations occur at the boundary.** “DNA exchange advantage” swarm agar plate of two non-kin wild type strains not carrying selective markers after 24h (left) and 48h (right) of incubation. No spontaneous mutants were observed to swarm into the area containing two antibiotics.

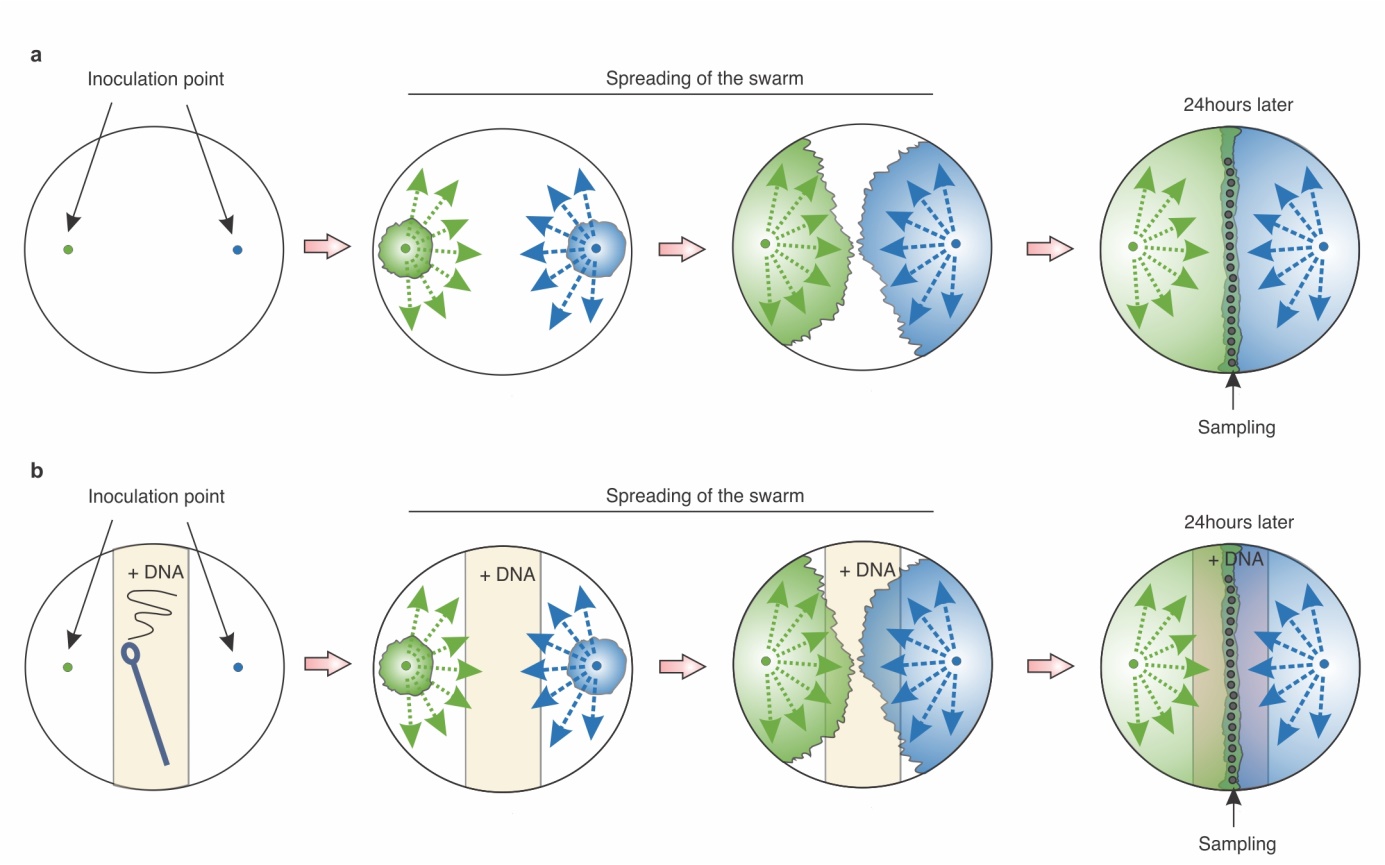

**Supplementary Fig. 9 |** **Experimental scheme for DNA exchange and DNA quantification experiment without and with added DNA. a,** Plates were inoculated with two non-kin strains (green and blue) and left for 24h incubation. At the meeting point where the boundary formed after 24h of incubation samples were taken (gray circles) and DNA exchange frequency and DNA concentration was determined as described in materials and methods. **b,** DNA (30 µg of DNA PS-216 ΔepsA-O (Tet^R^)) was added onto agar plate in a 1 cm wide stripe (yellow area) in the middle of the B medium agar plates. Samples were inoculated and sampled after 24h of incubation in the same manner as described above and materials and methods.

**Supplementary Results 1 | DNA exchange.** DNA exchange between non kin strains was significantly higher than DNA exchange between kin strains. For easier understanding we abbreviated strain names to ‘’Sp’’ (*amyE*::P43-*cfp* (Sp)) and ‘’Cm’’ (*sacA*::P43-*yfp* (Cm)). This section provides additional details about kin vs non-kin DNA exchange.

Kin DNA exchange between PS-13 (Cm) and PS-216 (Sp) was statistically similar to DNA exchange between isogenic strains PS-216 (Cm): ps-216 (Sp) (two tailed t-test, *p*=0.29) and to PS-13 (Cm):PS-13 (Sp) (two tailed t-test, *p*= 0.52), DNA exchange between PS-18 (Cm) and PS-216 (Sp) was statistically similar to DNA exchange between isogenic strains PS-18 (Cm): PS-18 (Sp) (two tailed t-test, *p*=0.17) and to PS-216 (Cm):PS-216 (Sp) (two tailed t-test, *p*= 0.52), DNA exchange between PS-68 (Cm) and PS-216 (Sp) was statistically similar to DNA exchange between isogenic strains PS-68 (Cm): PS-68 (Sp) (two tailed t-test, *p*=0.40) and to PS-216 (Cm):PS-216 (Sp) (two tailed t-test, *p*= 0.77). On the other hand, DNA exchange between non-kin strains PS-13 (Sp):PS-196 (Cm) was statistically different from DNA exchange between PS-13 (Cm):PS-13 (Sp) (two tailed t-test, *p*=0,002) and PS-196 (Cm):PS-196 (Sp) (two tailed t-test, *p*=0.0009), DNA exchange between non-kin strains PS-218(Cm):PS-216(Sp) was statistically different from DNA exchange between PS-218 (Cm):PS-218 (Sp) (two tailed t-test, *p*=0.003) and PS-216 (Cm): ps-216 (Sp) (two tailed t-test, *p*=0.005), DNA exchange between non-kin strains PS-196(Cm):PS-216(Sp) was statistically different from DNA exchange between PS-196 (Cm):PS-196 (Sp) (two tailed t-test, *p*=0.00002) and PS-216 (Cm): PS-216 (Sp) (two tailed t-test, *p*=0.00003). For *p*-values (two tailed t-test) between kin and non-kin DNA exchange combinations, see Supplementary Fig. 2.

When using swapped antibiotic markers, kin DNA exchange between PS-13 (Sp) and PS-216 (Cm) was statistically similar to DNA exchange between PS-216 (Sp): ps-216 (Cm) (two tailed t-test, *p*=0.89) and to PS-13 (Sp):PS-13 (Cm) (two tailed t-test, *p*= 0.44) (Supplementary Fig. 1a), DNA exchange between PS-18 (Sp) and PS-216 (Cm) was statistically similar to DNA exchange between PS-18 (Sp): PS-18 (Cm) (two tailed t-test, *p*=0.12) and to PS-216 (Sp):PS-216 (Cm) (two tailed t-test, *p*= 0.36), DNA exchange between PS-68 (Sp) and PS-216 (Cm) was statistically similar to DNA exchange between PS-216 (Sp):PS-216 (Cm) (two tailed t-test, *p*=0.48), however lower DNA exchange was observed between PS-68 (Sp): PS-68 (Cm) (two tailed t-test, *p*=0.02). On the other hand, DNA exchange between non-kin strains PS-13 (Cm):PS-196 (Sp) was statistically different from DNA exchange between PS-13 (Sp):PS-13 (Cm) (two tailed t-test, *p*=0,0012) and PS-196 (Sp):PS-196 (Cm) (two tailed t-test, *p*=0.0006), DNA exchange between non-kin strains PS-218(Sp):PS-216(Cm) was statistically different from DNA exchange between PS-218 (Sp):PS-218 (Cm) (two tailed t-test, *p*=0.019) and PS-216 (Sp): PS-216 (Cm) (two tailed t-test, *p*=0.0216), DNA exchange between non-kin strains PS-196(Sp):PS-216(Cm) was statistically different from DNA exchange between PS-196 (Sp):PS-196 (Cm) (two tailed t-test, *p*=0.048) and PS-216 (Sp): PS-216 (Cm) (two tailed t-test, *p*=0.050). For p values (two tailed t-test) between kin and non-kin DNA exchange combinations, Supplementary Fig. 2.

**Supplementary Results 2 | DNA concentrations** **at the boundary of wild type and nuclease mutant strains with the addition of exogenous DNA.** DNA concentration at the boundary of nuclease mutant non-kin strains (PS-216 *amyE*::p43-*cfp,* Δ*yhcR,* Δ*nuc*B and PS-196 *sacA*::p43-*yfp* Δ*yhcR*) was significantly higher than DNA concentration at the boundary of two wt strains (PS-216 and PS-196) without added DNA (*p*=0.001), and was found to be higher also when cca 30 µg of exogenously DNA PS-216 Δ*epsA-O* (Tet) was added (two tailed t-test, *p*=0.035). DNA concentration at the boundary of non-kin wt strain compared to DNA concentration at the boundary when the same wt strains were grown in the presence of cca 30 µg of exogenous DNA (PS-216 Δ*epsA-O* (Tet)) was statistically the same (two tailed t-test, *p*=0.90).

Nuclease mutants showed significantly higher DNA/CFU concentration in the boundary with or without exogenous DNA addition, compared to wt strains (two tailed t-test, without DNA *p*=0.0045, with DNA *p*=0.0053) (Fig. 4b) and boundary with nuclease mutants showed significantly higher concentrations of DNA/CFU if DNA was streaked on the agar plate before the experiment (two tailed t-test, *p*=0.012) (Fig. 4b) confirming that nucleases are the cause of lower DNA concentrations in the boundary of wt strains.

Nuclease mutants showed significantly higher DNA/mm^2^ concentration in the boundary with or without exogenous DNA addition, compared to wt strains (two tailed t-test, without DNA *p*=0.001, with DNA *p*=0.035) (Fig. 4b) and boundary with nuclease mutants showed higher concentrations of DNA/mm^2^ if DNA was streaked on the agar plate before the experiment, but the difference was not statistically significant due to oscillations in the measurements (two tailed t-test, *p*=0.159)

The amount of DNA in the boundary between wt strains and wt strain that grew with exogenously added DNA, was statistically similar for DNA/CFU (two tailed t-test, *p*=0.36) or per DNA/mm^2^ (two tailed t-test, *p*=0.902) at the boundary indicating that nucleases that are released are capable of degrading high amounts of exogenously added DNA.

**Supplementary Results 3 | DNA exchange between ∆*sigW* strains.** DNA uptake between non-kin ∆*sigW* strains (PS-216 and PS-196) compared to DNA uptake between wt non-kin strains decreased significantly for both strain pairs 1) PS-196 (Cm) ΔsigW : PS-216 (Sp) ΔsigW and for strain pair 2) PS-216 (Cm) ΔsigW : PS-196 (Sp) ΔsigW (p_1_=0.012, p_2_=0.053). Abbreviations ‘’Sp’’ and ‘’Cm’’ stand for (*sacA*::P43-*yfp* (Cm)) and (*amyE*::P43-*cfp* (Sp)), respectively. DNA uptake between non-kin PS-13:PS-196 and 216:218 ΔsigW mutants was under the detection limit, suggesting a dramatic decrease in the DNA transfer between the ΔsigW mutants.

The obtained lower DNA exchange frequency of pair 1 (PS-196 (Cm) ΔsigW : PS-216 (Sp) ΔsigW) resembled kin/self DNA exchange for PS-196 kin wt DNA exchange (PS-196 (Cm):PS-196 (Sp)(p_196_=0.35) and PS-216 wt DNA exchange (PS-216 (Sp): PS-216 (Cm) strain (p_216_=0.46). Likewise, the DNA exchange frequency of pair 2 (PS-216 (Cm) ΔsigW : PS-196 (Sp) ΔsigW) decreased to the kin level for both PS-216 and PS-196 kin wt DNA exchange frequency (p_216_=0.66, p_196_=0.36). The decrease in DNA exchange was detected in one strain pair with kin ∆*sigW* strains, namely PS-13 (p=0.02), and the increase in DNA exchange was detected in one strain pair with kin ∆*sigW* strain, namely PS-218 (p=0.01). Other kin ∆*sigW* mutants showed similar DNA exchange as wild type strains tested in kin combinations (p_216_=0.12, p_196_=0.71) (Fig. 4b).

**Supplementary Results 4 | DNA exchange advantage assay**. Pairs of non kin strains (PS-216 (*amyE*::P43-*cfp* (Sp) and PS-196 *sacA*::P43-*yfp* (Cm)) were tested for “DNA exchange advantage” on swarm agar plates and results presented in Fig. 6 and Supplementary Fig. 6 show that the transformants acquiring a resistance gene form non kin gain advantage and spread in the area containing two antibiotics. One half of the plate consisted of B medium and two non-kin strains were allowed to swarm towards each other without the presence of antibiotics (Fig. 6a). The other half of the plate was supplemented with two antibiotics (Sp and Cm) and spreading into the region containing both antibiotics was only allowed for DNA exchange transformants, carrying both antibiotic resistance genes (Sp and Cm) (Fig. 6a). The observed spreading area was sampled (Fig 6a and b, Supplementary Fig. 6) and inoculated onto agar plates containing Sp, Cm and Sp +Cm. The majority of the population isolated from the spread area consisted of PS-216 (*amyE*::P43-*cfp* (Sp))(Supplementary Fig. 7c), despite supplementation with two antibiotics, suggesting that the minority of transformants, carrying both Ab resistance genes helped the ancestor strain to spread by degrading antibiotics. Next, 50 randomly selected colonies obtained from Sp+Cm plates from 3 individual experiments (n_total_=150), were tested for *yfp* and *cfp* fluorescence with 100 and 98 % of all tested colonies growing on Sp and Cm (n=150) were positive for *cfp* and *yfp* fluorescence, respectively (Fig. 5d). When wild type strains were used in the “DNA exchange advantage” assay no spreading was observed after 24 or 48 hours of incubation demonstrating that spreading into area containing two antibiotics is the consequence of active marker gene acquisition and not spontaneous mutations at the boundary (Supplementary Fig. 8).
